# ATPase activity of *B. subtilis* RecA affects the dynamic formation of RecA filaments at DNA double strand breaks

**DOI:** 10.1101/2022.02.15.480544

**Authors:** Rogelio Hernández-Tamayo, Niklas Steube, Thomas Heimerl, Georg Hochberg, Peter L. Graumann

## Abstract

RecA plays a central role in DNA repair and is a main actor involved in homologous recombination (HR). *In vivo*, RecA forms filamentous structures termed “threads”, which are essential for HR, but whose nature is still ill defined. We show that RecA from *Bacillus subtilis* having lower ATP binding activity can still form nucleoprotein filaments *in vitro*, and still retains most of wild type RecA activity *in vivo*. Contrarily, loss of ATPase activity strongly reduces formation of nucleoprotein filaments *in vitro*, and effectivity to repair double strand breaks (DSBs) *in vivo*. While lowered ATP-binding activity only moderately affected RecA dynamics, loss of ATPase activity lead to a large reduction of the formation of threads, as well as of their dynamic changes observed in a seconds-scale. Single molecule tracking of RecA revealed incorporation of freely diffusing and non-specifically DNA-bound molecules into filaments upon induction of a single DSB. This change of dynamics was highly perturbed in the absence of ATPase activity, revealing that filamentous forms of RecA as well as their dynamics depend on ATPase activity. Our data suggest that RecA/ssDNA filaments change in subcellular localization and length involving ATP-driven homology search.

## INTRODUCTION

All cells possess an intricately regulated response to DNA damage. Bacteria have evolved an extensive regulatory network called the SOS response to control the synthesis of factors that protect and repair the genome. Processes co-ordinately regulated within the SOS response include error-free DNA repair, error-prone lesion bypass, inhibition of cell division, and homologous recombination (HR).

HR is a mechanism used by all organisms to repair DNA lesions such as double-stranded breaks (DSBs) and collapsed replication forks. DSBs can be repaired through two principally different processes: error-prone direct end joining, or recombination with the intact DNA duplex of the sister chromosome (1). To carry out its central function in HR, RecA must assemble into helical nucleoprotein filaments on the ssDNA tracts formed at sites of DSBs or stalled replication forks. Although RecA binds rapidly to ssDNA in purified systems, its ability to bind tracts of ssDNA *in vivo* requires one of two pathways that allow RecA to outcompete the high affinity ssDNA binding protein SSB. Initially, during presynapsis, DNA ends are primed for loading of the central homologous recombination protein RecA (Rad51 in eukaryotes) by the MRX (Mre11–Rad50–Xrs2) complex in eukaryotic cells or by RecBCD, RecN, RecF, RecO, and RecR proteins in bacteria (1–5). During synapsis, RecA sets up strand exchange by introducing a single DNA strand from the break site into the intact homologous sister chromosome, and vice versa. Finally, during post synapsis, RecA-mediated three-way junctions are converted into true crossovers (or Holiday junctions) through the action of proteins such as the RecG helicase, and branch migration and resolution of Holiday junctions are achieved through the action of the RuvABC complex in bacteria (6). RecA is a DNA-dependent ATPase (7–9). The ATP bound form of *Escherichia coli* RecA has a higher affinity for ssDNA and dsDNA than does the ADP-bound form (10). Thus, ATP hydrolysis converts a high affinity DNA binding form, RecA-ATP, to a low affinity form RecA-ADP, thereby supporting an ATP hydrolysis-dependent dynamic cycle of DNA binding and dissociation.

This difference in affinity, combined with the protein’s DNA dependent ATPase activity, results in a dynamic DNA binding cycle: RecA-ATP cooperatively binds DNA to form helical nucleoprotein filaments; ATP hydrolysis throughout the filament converts RecA-ATP to RecA-ADP; and RecA-ADP protomers dissociate from DNA. Filament extension on ssDNA is less cooperative, possibly resulting in the formation of short filament patches as observed for RAD51 (7,10). The 3’end is typically favoured for addition of RecA monomers and disassembly occurs from the 5’ end resulting in a net 5’ to 3’ assembly direction on ssDNA (11,12). Due to the 5’ to 3’ directionality, the 3’ end is more likely to be covered with RecA resulting in more efficient pairing reactions at the 3’ end *in vitro* (13,14). Resolved structures of RecA complexed with ssDNA indicate that ATP binding and ATP hydrolysis indeed mediate the binding and release of RecA from DNA through allosteric coupling (15). ATP hydrolysis is, however, not the only driving factor for dissociation, as the mechanical interactions of a recombinase filament with its stretched DNA substrates also facilitate disassembly (16). Although RecA is a DNA-dependent ATPase, its homology search and strand exchange activities are largely independent of its ATPase activity (17).

Proteins that have ATP hydrolytic activity typically have two well-defined motifs called the Walker A and B motifs (18). Researchers wishing to test the importance of ATP binding and hydrolysis in biological roles for these proteins create two types of mutations in the Walker A or P-loop motif. Both mutate the highly conserved lysine residue. A conservative change of the lysine residue to an arginine in many cases creates a protein that can still bind ATP but is no longer able to hydrolyse it (19–21). The second type replaces the lysine with an alanine. Here, the protein is commonly found to have greatly reduced ability to bind ATP (22–25). Despite RecA having been analysed in depth in the Gram-positive model bacterium *Bacillus subtilis in vivo* and *in vitro*, loss of ATP binding or of ATPase activity have not yet been tested *in vivo*.

Interestingly, when DNA modifications are induced that lead to the generation of double strand breaks, or if defined DSBs are induced in the chromosome of bacteria, RecA assembles into filament-like structures that have been termed “threads” (26). It has been suggested that filamentous RecA structures are ssDNA RecA filaments that guide ssDNA from the break site towards the homologous DNA region that is generally located in the other cell half in *B. subtilis* cells (26), or to form protein filaments that guide a chromosome site containing a DSB towards the non-broken sister site in the other cell half (27). Because in general, bacteria rapidly segregate replicated regions on the chromosome into opposite cells half, sister loci are separate from each other, and are moved together in *E. coli* or *Caulobacter crescentus* cells, when a DSB is induced in one site (27–29). RecA has been proposed to mediate repairing of separated sister loci, by providing a track for e.g. motor proteins moving DSB sites (27), or by providing directionality for the search of the sister site (29) that is found due to sufficient sequence homology. In *B. subtilis* cells grown to the state of competence, DNA is taken up at a single cell pole. From this site, RecA forms threads that have been shown to be essential for homologous recombination during transformation with externally acquired DNA (30). Filaments have been shown to be about 60 nm thin (29), and to remodel within a time frame of minutes in several studies (26,27,29,30), possibly performing a motor-like function. Especially, how filament dynamics are driven is still unclear, as well as their mode of action.

In this study, we addressed the question if loss of ATP binding or ATPase activity affect properties of RecA *in vitro*, or thread dynamics *in vivo*. *RecA*_K70R_ and *recA*_K70A_ alleles were placed into the *recA* locus, and translational fusions to mVenus or sfGFP were generated. These strains were assayed for their ability to repair and recombine DNA, and to induce a DSB-response by forming dynamic RecA threads on the DNA. Dynamics of RecA were assayed by superresolution (SIM) light microscopy and at single molecule level *in vivo*, yielding insight into the role of RecA ATPase activity for nucleating RecA filaments. In our work, we developed a probe that specifically visualizes and quantifies RecA structures on DNA a using single molecule tracking approach and utilize it to provide a detailed timeline of RecA structural organization in living cells after DSBs. We show that biochemical properties of *B. subtilis* RecA differ from those reported for *E. coli* RecA with respect to changes in ATP binding or ATPase activity, and that the latter is crucial for filament dynamics *in vivo*. This finding has important implications for the conclusions that can be drawn from filament dynamics observed in live cell imaging.

## RESULTS

### Characterization of Walker A mutations in RecA *in vitro*

RecA ATPase activity has been studied extensively *in vitro* (17,31), but its requirements for recombination, DNA repair, and DSBs induction in the Gram-positive model bacterium *B. subtilis* have not yet been analysed *in vivo*. Wild type RecA requires ATP binding for efficient loading onto ssDNA, and shows ATPase activity during DNA strand exchange, and also during unloading from ssDNA filaments (32). A major aspect of our work was to determine if the formation of dynamic thread structures requires ATP binding or ATP hydrolysis *in vivo*. To this end, we purified wild type RecA, and two Walker A mutant forms expected to abolish ATP binding (K70A) or to allow for ATP binding but prevent ATPase activity (K70R) as C-terminal hexa-histidine fusion proteins, via nickel-NTA affinity chromatography, and in a second step via size exclusion chromatography. Fig. 1A shows that all three variants eluted in a single peak corresponding to RecA monomers. However high salt conditions (500 mM NaCl) that were required to remove residual, bound DNA from the overexpression host, may drive RecA into its monomeric form (see below). Fig. 1B shows that all variants were obtained in high purity. For all three variants, we did not observe any ATPase activity in the absence of ssDNA (Salmon Sperm ssDNA), but surprisingly, we found lower, but substantial activity for the RecA_K70A_ mutant, while there was no detectable activity for RecA_K70R_ (Fig. 1C). This is due to residual ATP binding for RecA_K70A_, which was about three-fold lower compared to that of wild type RecA, in the presence of ssDNA. RecA_K70R_ mutant also showed about three-fold reduced ATP-binding activity, while in the absence of ssDNA, there was no binding activity, which for wild type RecA was also strongly reduced (Fig. 1D).

**Figure 1.**
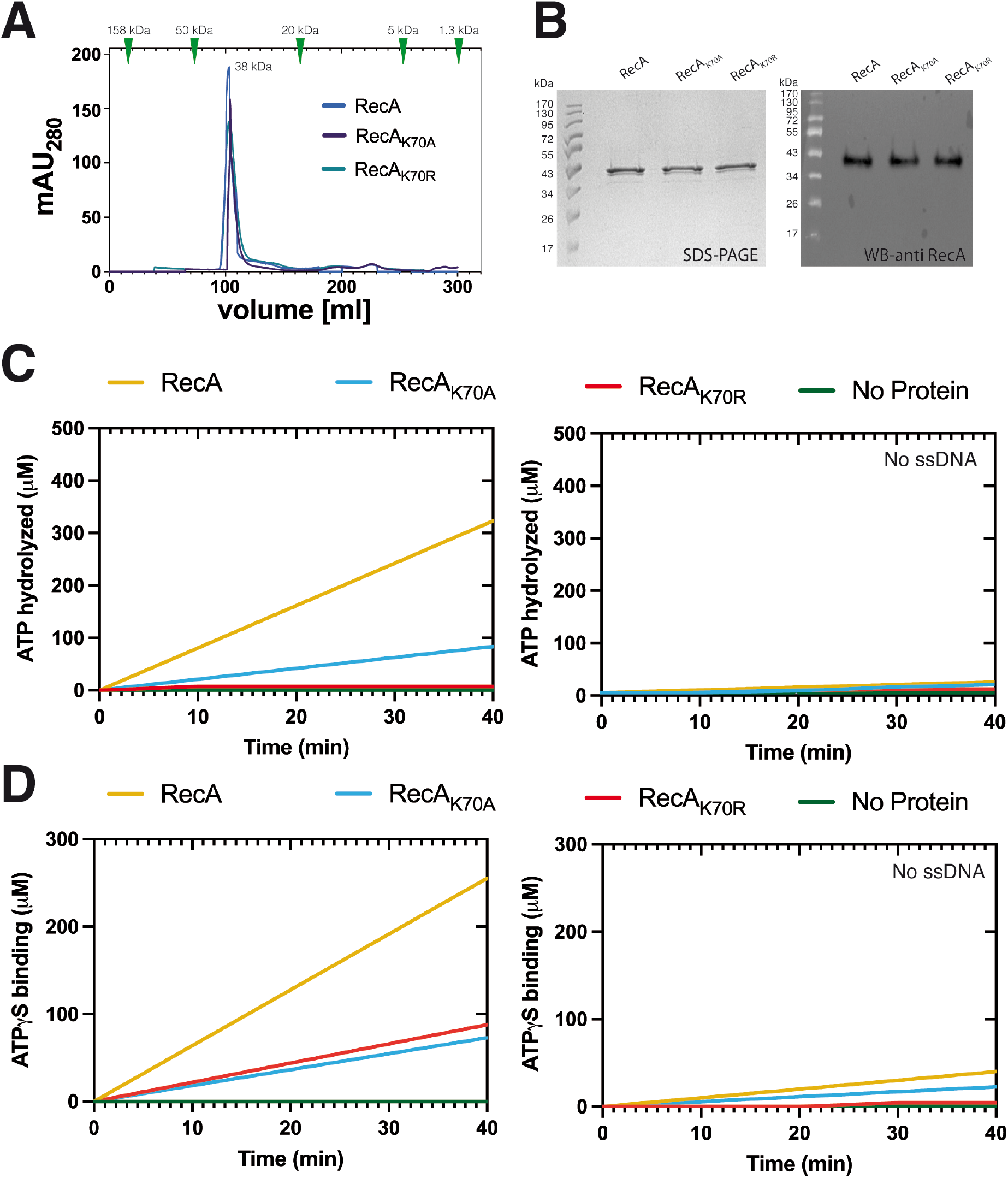
Biochemical characterization of RecA Walker A mutants. A) Gel filtration (GF) of His Tag-RecA_WT_ and mutants after Nickel-Sepharose affinity chromatography and ensuing size-exclusion chromatography. Size standards are shown with triangles in the upper part of the chromatogram. B) SDS-PAGE and Western blot (WB) of fractions corresponding to elution fractions in GF fractions, for WB, proteins were probed using a 1:5000 dilution (rabbit-α-RecA) and secondary goat-α-rabbit-antibody (1:10000 dilution). C) ATP hydrolysis assay as described in Material and Methods. D) Kinetic analysis of ATP binding assay as described in Material and Methods.

Thus, while the lysine to arginine substitution behaved roughly to our expectation, losing ATPase activity but not completely ATP binding, the alanine exchange did not result in a complete loss of ATP binding, but to a strong reduction and thus also to decreased ATPase activity. These findings put us into the situation of addressing the question if loss of ATPase activity leads to changes in RecA activity and/or formation of threads during DNA repair.

### Loss of ATPase activity affects nucleofilament formation *in vitro*

We wished to analyse the effects of Walker A mutations in *B. subtilis* RecA on the formation of nucleofilaments *in vitro*. We therefore analysed RecA bound to ssDNA of various length (main length 580 to 800 nt) by electron microscopy (EM) and determined the size of nucleofilaments with oligonucleotides of 32 nt using mass photometry (MP). Fig. 2A shows that ssDNA by itself did not yield substantial MP signals, while RecA and the two mutant versions each showed a well-defined peak around 80 kDa, consistent with the theoretical mass of 76 kDa for dimers. If monomers are populated in solution, they may not have been observable in our MP experiments, because their mass (38 kDa) is close to the detection limit of the instrument. Our MP measurements do not agree with our SEC experiments (Fig. 1A), in which we only observed monomers for all three variants. This is likely a result of the much higher salt concentrations used in the SEC experiment compared to the MP measurement (500 mM NaCl and 137 mM, respectively), wherefore we favour the view that under physiological conditions within cells, RecA forms dimers.

**Figure 2.**
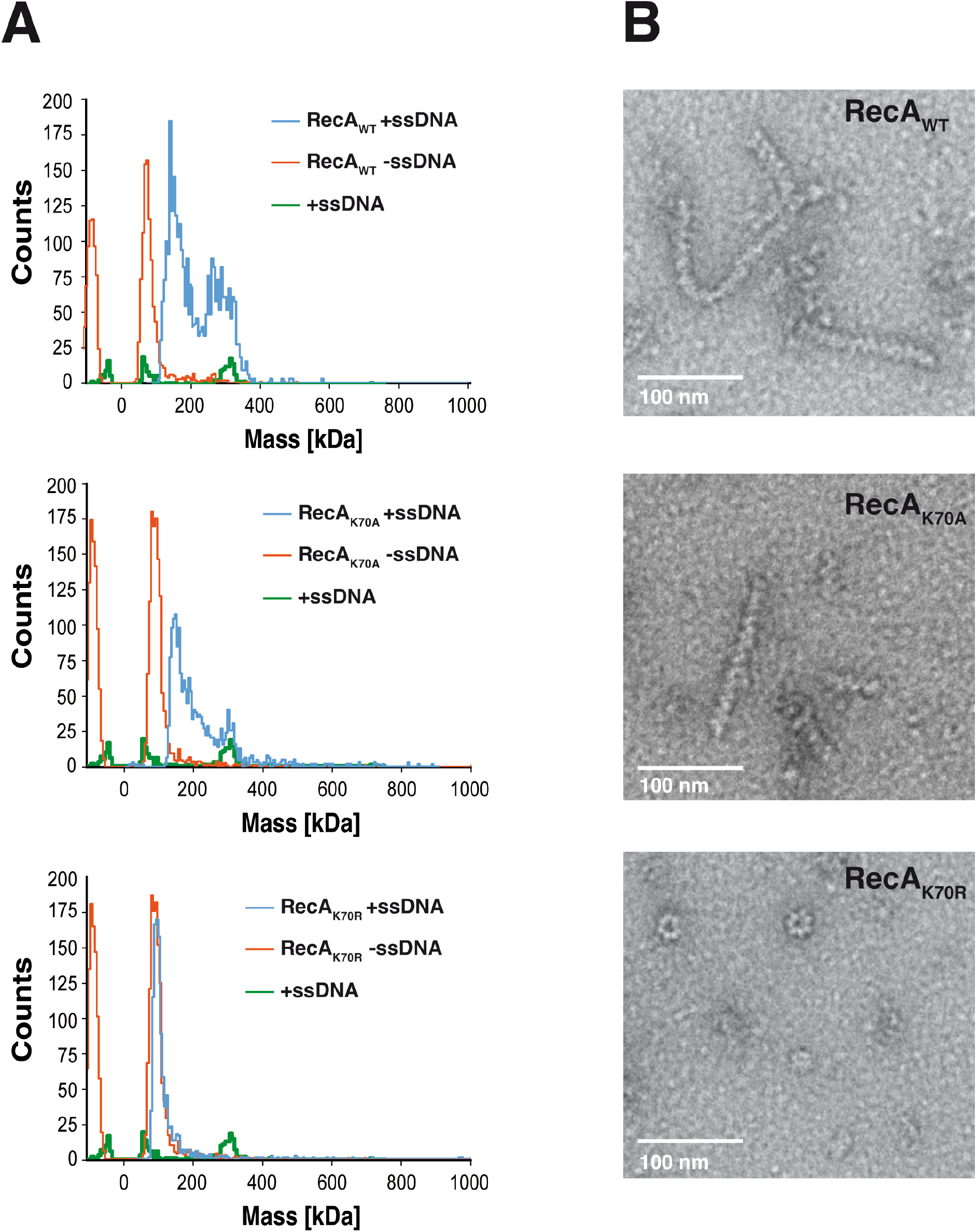
Filament formation of RecA and ATPase mutants in mass photometry and electron microscopy. A) Molecular mass distribution histograms of RecA_WT_, RecA_K70A_ or Reca_K70R_ alone or incubated with ssDNA and ATP. B) Electron micrographs show filament formation by negative staining of ssDNA and wild type RecA or Walker A mutants.

When incubated together with ssDNA, mass spectra of RecA showed two peaks, a well-defined peak at about 160 kDa, and a broad peak between 250 and 400 kDa (Fig. 2A). Both peaks are larger than dimers, indicating that they represent DNA-bound species. We could not determine the exact stoichiometry implied by these peaks, because our measurements were calibrated with protein standards, which are inaccurate for DNA-containing complexes (33). In addition, we could not determine a pseudo- (protein calibrated) mass of our free ssDNA oligo, because the oligo yielded no appreciable scattering on their own, even when using poly-L-lysine coated slides. Regardless, our MP measurements of Rec A WT clearly show ssDNA binding at nanomolar concentrations. We could verify the presence of nucleoprotein filaments by EM (Fig. 2B). Helicity of NPFs is clearly visible, in agreement with what was shown for *E. coli* RecA. For RecA_K70A_, we also observe DNA binding by MP and EM analyses, however revealing somewhat reduced formation of RecA-ssDNA filaments.

MP spectra of the K70R mutant showed a single peak at around 80 kDa in the presence of ssDNA (Fig. 2A), consistent with free dimers, and only very few polymeric structures were visible by EM. Interestingly, those corresponded to ring-shaped heptamers (with dimers not being well visible), indicating that there is a low propensity for the formation of polymers with a different architecture than wild type polymers (Fig S1). We could not observe these heptamers in our mass photometry assays, likely because of the much lower concentrations used in MP experiments. In *E. coli*, RecA_K70R_ has virtually no ATP hydrolytic activity, but can bind ATP, and ssDNA (19). Thus, *B. subtilis* RecA lacking ATPase activity shows a strikingly different effect of no longer binding to ssDNA. Together, our results show that ATP hydrolysis is essential for the formation of RecA-DNA filaments *in vitro*.

### Loss of ATPase activity affects nucleofilament formation *in vitro*

While RecA is highly conserved in bacteria, its activity and regulation can be quite different between phyla. We wished to analyse the effect of Walker A mutations in *B. subtilis* RecA on the formation of nucleofilaments *in vitro*. We therefore analysed RecA bound to ssDNA of various length by electron microscopy (EM) and determined the size of nucleofilaments using mass spectroscopy. Fig. 2A shows that ssDNA by itself did not yield substantial signals, while RecA and the two mutant versions showed a peak just below 100 kDa, likely representing dimers (76 kDa). For mass spectroscopy, low-salt conditions were employed, allowing for dimer formation, wherefore we favour the view that RecA exists as dimers under native conditions. The same behaviour of RecA and the two mutant versions indicate that dimer formation does not depend on ATP binding, or else we would have expected to see two peaks (the system can accurately determine protein size from 50 kDa up).

Interestingly, when incubated together with ssDNA, RecA showed two peaks, a well-defined peak at about 160 kDa, and a broad peak between 250 and 400 kDa (Fig. 2A). The latter likely represents RecA nucleoprotein filaments (NPFs), which was verified by EM (Fig. 1B). Helicity of NPFs is clearly visible, in agreement with what was shown for *E. coli* RecA. For RecA_K70A_, the fraction of NPFs was somewhat reduced, but EM analyses clearly showed formation of ecA7ssDNA filaments. However, K70R mutant RecA showed a single peak close to 100 kDa (Fig. 2A), characteristic of protein-only structures (Fig. 2A), and only very few polymeric structures were visible by EM. Interestingly, those corresponded to ring-shaped heptamers (with dimers not being well visible, Fig. S1), indicating that there is a low propensity for the formation of polymers with a different architecture than wild type polymers. In *E. coli*, RecA_K70R_ has virtually no ATP hydrolytic activity, but can bind ATP, and ssDNA (19). Thus, *B. subtilis* RecA lacking ATPase activity shows a strikingly different effect of no longer binding to ssDNA *in vitro*.

### Effects of Walker A mutations on the functionality of RecA

We next moved our investigations to *in vivo* conditions. *RecA*_K70A_ and *recA*_K70R_ alleles were transferred into the chromosome of *B. subtilis*, under the control of the original *recA* promoter, generating a merodiploid strain in which the original *recA* gene is driven by the xylose promoter, and is not induced during normal growth. As mentioned above, *E. coli* RecA_K70R_ is dysfunctional, and is has been shown to display a reduced SOS response activity compared to wild type RecA (34). We tested for a defect in DNA repair via recombination by using an established system in which the HO endonuclease gene is integrated into the *amyE* locus on the chromosome, driven by the inducible xylose promoter, and the corresponding cut site close to the origin of replication. Expression of HO endonuclease was induced for a different length of time, followed by plating on plates lacking inducer, generating a DNA cut close to the origin regions of replication; this leads to cuts in about two-thirds of chromosomes in an exponentially growing *B. subtilis* culture (26,35). As most cells contain separated origin regions for most of the cell cycle (36), some cells will contain no cut, some a single cut, and some cuts in both origin regions. In wild type cells, colony formation stayed roughly constant for the first 75 minutes (Fig. 3A). In the RecA_K70A_ strain, there was a minor loss of viability, while DSBs repair was strongly affected in the RecA_K70R_ strain. However, the latter retained some activity, in contrast to a *recA* mutant strain, which failed almost completely to deals with DSBs (Fig. 3A). Mutant proteins were expressed at wild type levels (Fig. 3B). Residual repair activity in the ATPase mutant strain may be due to expression of the wild type *recA* gene for the length of HO induction, as a side effect from induction by xylose.

**Figure 3.**
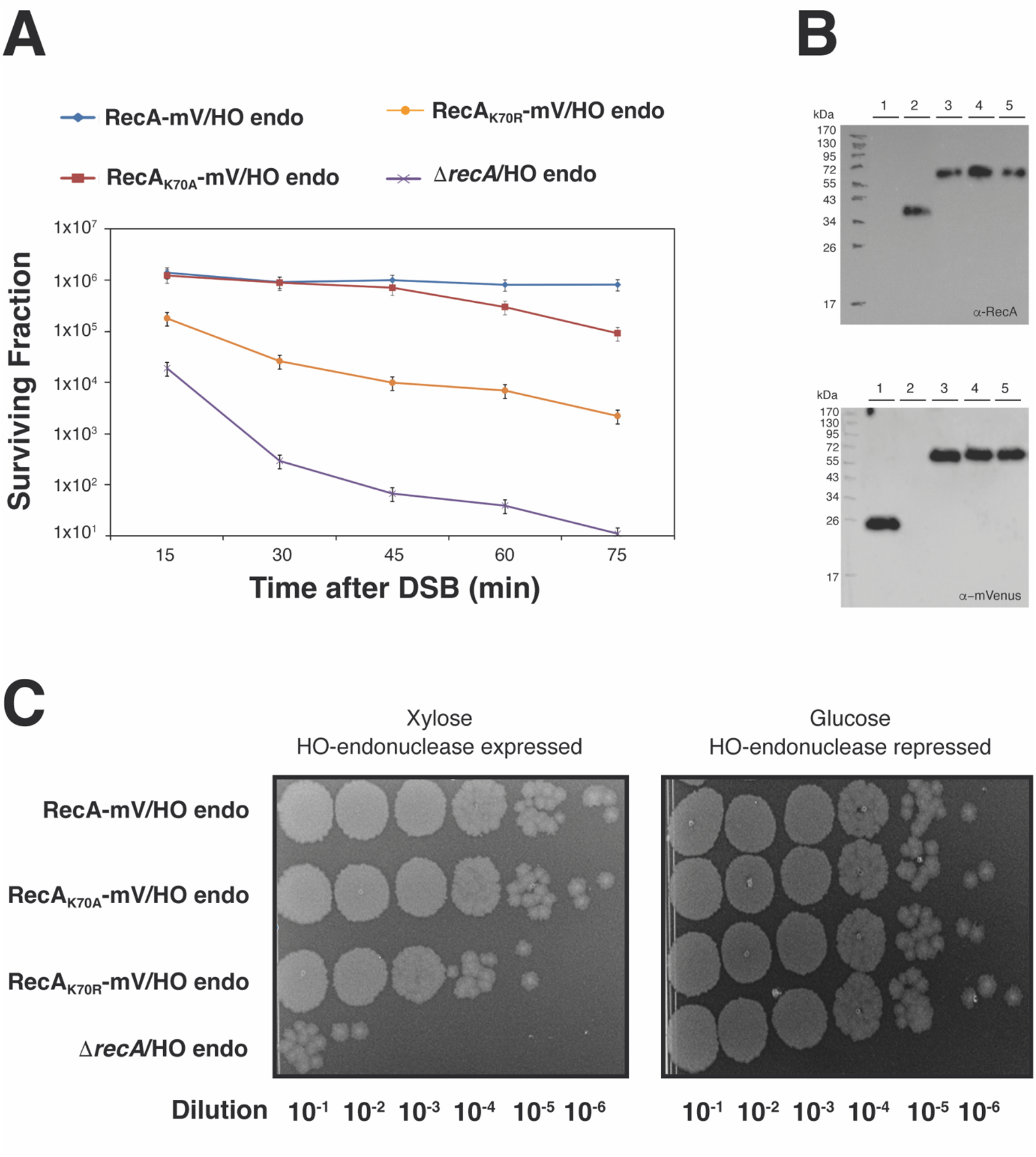
Survival assays and effects of RecA ATP-binding or ATPase mutants on the functionality of RecA. A) Survival curves of strain RecA-mV/HO endo, RecA_K70A_-mV/HO endo, RecA_K70R_-mV/HO endo and Δ*recA*/HO endo, after induction of the DSB. All strains were plated in duplicate, in three independent experiments. B) Western blots of fluorescent protein fusions. Shown are Western blots of from whole cell lysates of 1) *E. coli* expressing mVenus, 2) *B. subtilis* BG214, or 3) RecA-mV/HO endo, 4) RecA_K70A_-mV/HO endo and 5) RecA_K70R_-mV/HO endo expressing cells. Proteins were probed using a 1:500 dilution (rabbit-α-GFP) and secondary goat-α-rabbit-antibody (1:10000 dilution) or using a 1:5000 rabbit-α-RecA dilution and secondary goat-α-rabbit-antibody (1:10000 dilution. Cells were harvested in exponential phase at OD_600_ 0.5 – 0.7 prior to analysis. C) Spot assays. Cultures of RecA-mV/HO endo, RecA_K70A_-mV/HO endo, RecA_K70R_-mV/HO endo and Δ*recA*/HO endo, before and after DSB-induction.

The same effects were qualitatively found using spotting assays after HO endonuclease induction (Fig. 3C). These data show that reduced ATP binding activity does not severely impair RecA function *in vivo*, unlike loss of ATPase activity, which severely affects RecA activity.

### ATPase activity is essential for the dynamics of RecA threads *in vivo*

GFP-RecA has been shown to first form a fluorescent focus at a site of a DSB, which then extends into thread structures that remodel on a time scale of few minutes (26). A major goal in this study was to investigate if the dynamics of RecA filaments depend on its ATPase cycle. We therefore used super resolution fluorescence microscopy, to obtain a better resolution of filamentous structures, and an even higher temporal resolution than earlier. We used structured illumination microscopy (SIM) on a fully functional RecA-mNeonGreen fusion (37) and acquired images every 20 seconds, using 500 ms exposure time. Fig. 4A shows that RecA threads changed in their subcellular arrangement between 20 s intervals (movie S1). Foci or threads can appear between 20 second intervals, and gain in fluorescence intensity. Threads frequently extend until they split into several entities, which can shift in their subcellular position. This behaviour is very similar to what has been shown before using a lower temporal resolution. RecA dynamics were quite similar for RecA_K70A_, where we also observed fluorescent foci and threads of high fluorescence intensity (Fig. 4B, movie S2). In striking contrast, RecA_K70R_ formed structures with much less intensity, and these did not show dynamics like the wild type protein (Fig. 4C, movie S3). To better visualize these finding, we generated demographs, in which actual signal intensities were plotted against cell length, showing that the alanine mutation somewhat reduced signal intensities, while the arginine mutation drastically lowered focus/filament formation (Fig. 4D). Kymographs in Fig. 4E show examples of cells, in which signal intensities of wild type RecA frequently changed their subcellular positioning, while this was strongly reduced in ATPase mutant RecA. These finding reveal that loss of ATPase activity strongly reduces assembly of RecA into foci (i.e., assembling at DSBs) and into threads, while reduction of ATP binding has a noticeable but less pronounced effect on thread formation and their dynamics.

**Figure 4.**
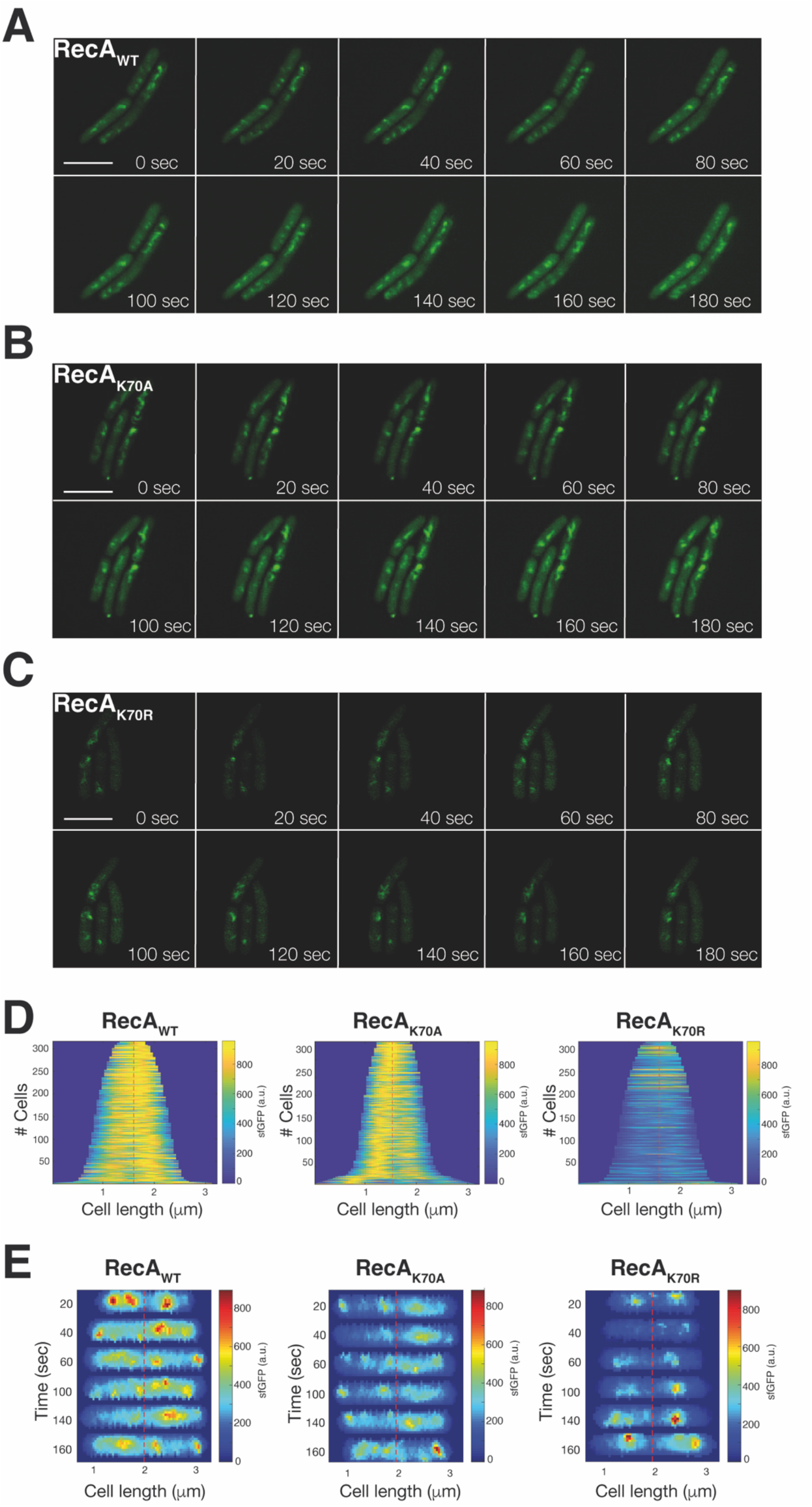
Structured illumination microscopy imaging of RecA thread formation in *B. subtilis* during double strand break repair. A-C) Montage of movies (shown in supplementary material) with SIM reconstruction for RecA_WT_-sfGFP, RecA_K70A_-sfGFP and RecA_K70R_-sfGFP. Site specific DSBs were induced by the addition of xylose to exponentially growing cells. Z-stacks resulting from the sfGFP channel were merged and projected into tomographic representations. 20 ms acquisition time, scale bar 2 μm. D) Demographs of RecA_WT_, RecA_K70A_ and RecA_K70R_ *B. subtilis* cells, showing the localization of RecA-sfGFP to the central regions. Cells were aligned and ordered according to size. The fluorescence profiles represent the mean fluorescence values along the medial axis after background subtraction and normalization such that the maximum fluorescence of each cell is equal. E) Kymographs show examples of cells, in which signal intensities of RecA_WT_-sfGFP or of RecA_K70A_-sfGFP, but not of RecA_K70R_-sfGFP, frequently changed their subcellular positioning.

### *In vivo* dynamics of RecA at a single molecule level

Dynamics of RecA have been visualized at single molecule level in *in vitro* experiments, but to our knowledge, there are no experiments of single molecule motion *in vivo*. We also used the functional RecA-mVenus (“mV”) fusion to track single molecule dynamics during exponential growth, or after induction of DSBs. This was done using 20 ms stream acquisition, and we discarded tracks of less than 5 steps in order to avoid bias based on very short tracking events. Of note, we have shown that use of UV or blue light excitation slows down growth of *B. subtilis* cells or other bacteria, but not green light excitation (38), suggesting that use of 514 nm laser illumination does not induce harm to cells that might alter their physiological state. Mean squared displacement analyses showed that wild type RecA and RecA_K70A_ had similar diffusion rates, while RecA_K70R_ moved much slower through the cells (Fig. 5A). After induction of a DSB via induction of HO endonuclease, wt RecA showed much decreased mobility (Fig. 5A). This decrease was less pronounced for ATP binding-mutant RecA_K70A_, while ATPase mutant RecA_K70R_ showed the opposite behaviour, with molecules becoming much more mobile. To better interpret these drastic effects, we employed squared displacement analyses to determine if monitored tracks might fall into different categories of mobility. The visualization of the results is shown in Fig. 5C in the form of jump distances, where the probability of different lengths of squared displacements (representing the number of certain lengths of motion done by molecules) is plotted. A single population of molecules would yield a single continuous probability distribution, or Rayleigh distribution. To explain all RecA tracks in a satisfactory manner, we needed to fit the data with three Rayleigh distributions (Fig. S2), resulting in an R^2^ value of 0.99. Using this fitting, we found a slow population of 12% (Fig. 5B, red curve in 5C), a medium-mobile population of 47% (black curse, Fig. 5C), and a high mobility fraction of 41% (blue curve, Fig. 5C). Taken together, the three Rayleigh fits result in the fit indicated by the grey dotted line, which fits the data very well (Fig. S2). The different states of mobility can be most easily explained assuming freely diffusive RecA for the high-mobility population, RecA binding non-specifically to dsDNA in the chromosome (medium mobility), and RecA being tightly bound ssDNA occurring in some cells due to spontaneously arising DNA damage. Fig. 5C shows that track length for RecA_K70A_ is shifted to slightly larger values, resulting in an increase of likely freely diffusive RecA molecules at the expense of medium-mobile RecA. Strikingly, RecA_K70R_ mutant molecules showed strongly down-shifted track lengths, in agreement with the much lower MSD value (Fig. 5A). ATPase mutant RecA showed more than 40% of molecules in the slow-mobile state, at the expense of medium and fast-mobile molecules (Fig. 5B and C). Because lack of ATPase activity is expected to abolish RecA activity in active strand exchange and based on abnormal interaction of mutant RecA with ssDNA (Fig. 2B), we favour the view that mutant RecA_K70R_ forms non-productive interactions with ssDNA and/or aggregates that lead to low mobility. In any event, our analyses show that loss of ATPase activity strongly affects RecA dynamics even during exponential growth.

**Figure 5.**
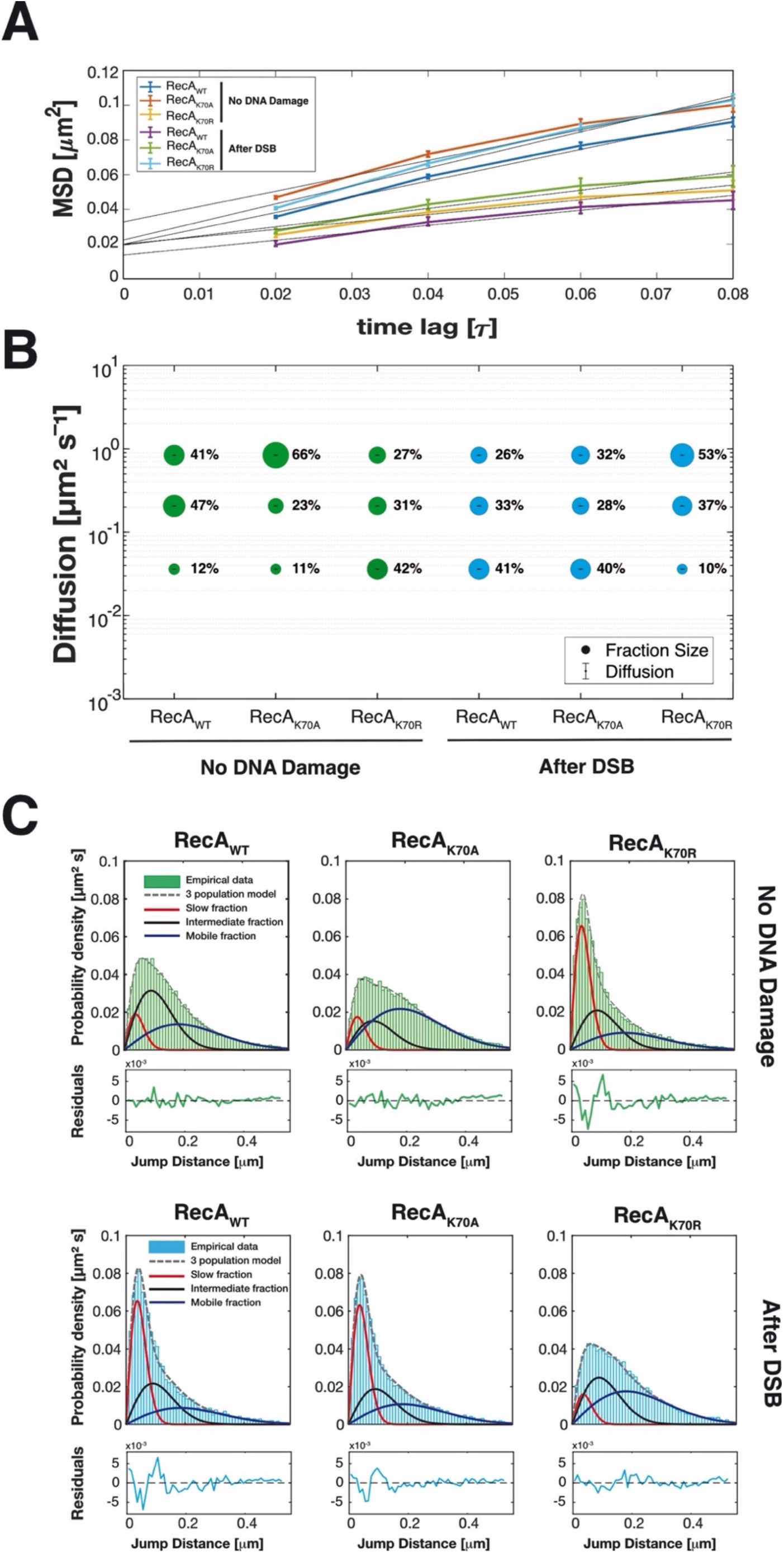
Analyses of single-molecule dynamics of RecA_WT_-mV, RecA_K70A_-mV and RecA_K70R_-mV expressed in *B. subtilis* cells during exponential growth. A) Mean squared displacement (MSD) analyses of RecA_WT_-mV, RecA_K70A_-mV, and Reca_K70R_-mV, showing different overall diffusion constants. MSD assumes an overall diffusion regardless of sub-diffusion. B) Bubble plot, derived from squared displacement analyses (SQD) shows the size of the fractions (proportional to the area) and corresponding diffusion coefficients in cells with no DNA damage and after DSB-induction. C) Jump distance analysis shows probability of displacements, different coloured solid lines represent the subpopulations; dotted lines represent the sum of the subpopulations.

### Loss of ATPase activity abolishes the response of RecA dynamics to the induction of double strand breaks

After induction of HO endonuclease, inducing a single DSB [or two, the system employed leads to cuts in about 75% of all chromosome loci (26)], population sizes for wild type RecA changed strongly, increasing 3.5 fold from 12 to 41%, while medium and fast-mobile fractions decreased accordingly (Fig. 5C). The visible formation of thread-structures (Fig. 4) supports the idea that the slow-mobile fraction of RecA molecules are those engaged in filament formation, suggesting that almost 50% of RecA is converted into its filamentous form. A very similar trend could be seen for ATP binding-mutant RecA, while ATPase mutant RecA showed completely opposing changes, in that the slow-mobile fraction became depleted, such that molecules had much higher mobility than before damage induction (Fig. 5B and C). Based on our assumption that the slow-mobile fraction refers to ssDNA-bound RecA, these experiments support the finding that ATPase mutant RecA fails to form dynamic filamentous structures *in vivo* (Fig. 4), and strongly malfunctions (Fig. 3).

In order to visualize changes in single molecule dynamics in 2D, all tracks were projected into a medium-sized cell of 3 x 1 μm (note that *B. subtilis* is about 0.8 μm wide), and were sorted into those that show very little motion for more than XY time intervals (“confined” motion, determined from three times the localization error), illustrated by red tracks in Fig. 6, those that show free diffusion (blue tracks), and those that show transitions between confined and free motion (green tracks, “transitions”). During exponential phase, wild type RecA reveals confined motion on the central part of cells containing the nucleoids, and free diffusion throughout cells (Fig. 6). After induction of DSBs, confined motion becomes more focussed on the nucleoids, and transition events are visibly reduced, indicating that many molecules have been tightly incorporated into putative RecA/ssDNA filaments. RecA_K70A_ does not feature strong changes in the pattern of motion, the increase in confinement seen from SQD analyses is less visually apparent. Note that slow-mobile molecules will largely agree with molecules showing confined motion but are not identical with this fraction. Conversely, RecA_K70A_ revealed opposing changes, confinement was visibly highly reduced following induction of DSBs, while exponentially growing cells looks almost indistinguishable from wild type cells during DSB repair. Possibly, lack of ATPase activity leads mutant RecA stuck in filaments that cannot be depolymerized, such that infrequent repair events during growth fail to be resolved.

**Figure 6.**
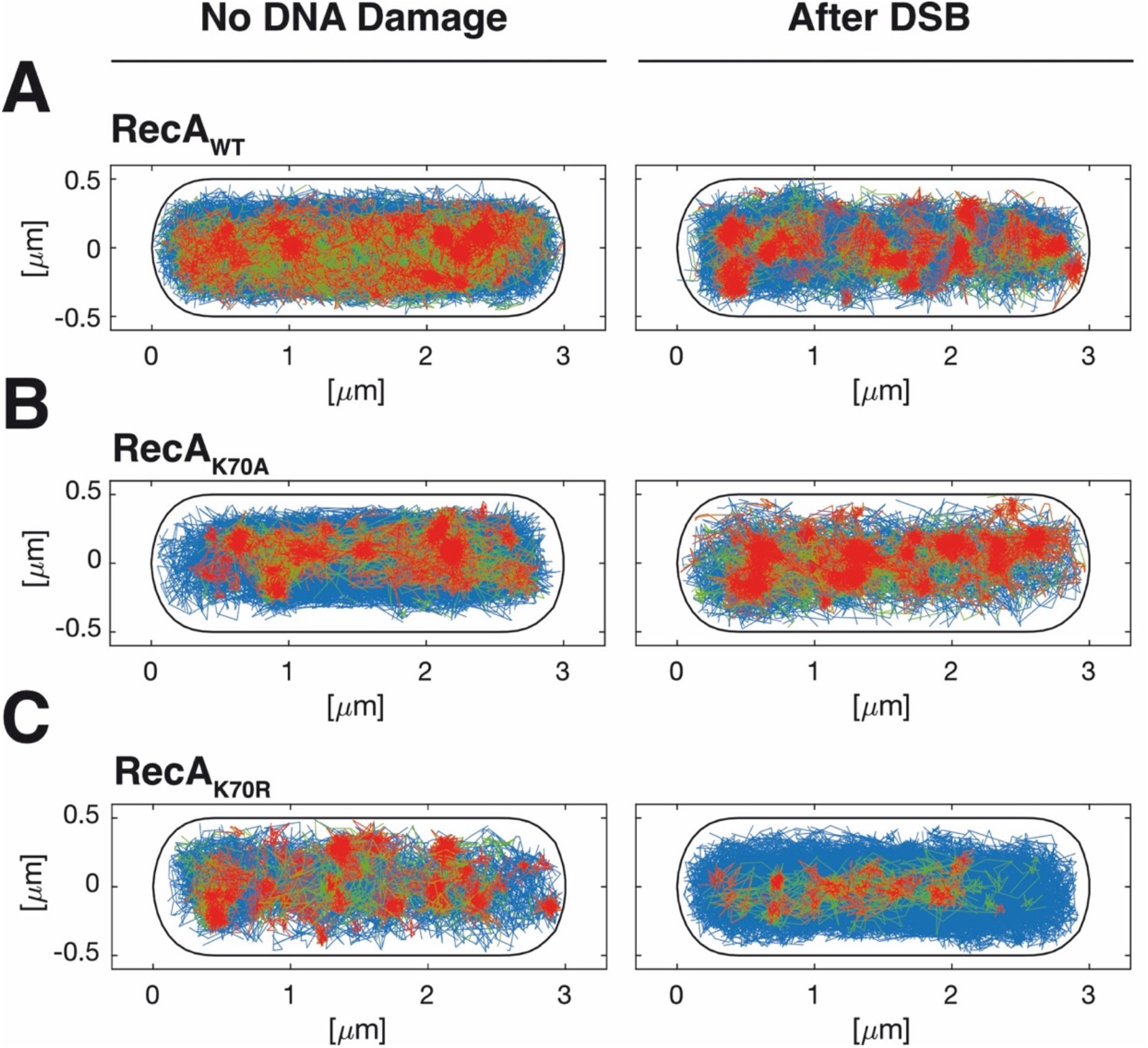
Confined motion of RecA molecules is restricted to nucleoid areas in the cell. **A-C)** Confinement maps are an algorithm of SMTracker program, and are shown for RecA_WT_-mV, RecA_K70A_-mV, and Reca_K70R_-mV with no DNA damage and after induction of a DSB. A trajectory is considered to present confinement (red) when it has a dwell event (molecules staying within a radius of 120 nm) for at least 5 time points. Molecules changing between confinement and mobility are termed “transition” (mixed behaviour), shown in green, and freely diffusive molecules lacking considerable parts of confinement are shown in blue.

## DISCUSSION

RecA and Rad51 are the central players in homologous recombination, which in turn is at the heart of repair of DNA damage such as double strand breaks in a chromosome, transformation with foreign DNA, or restart of stalled replication forks (39–42). While Rad51 can generally rely on spatially paired sister chromosomes during DSB repair in S-phase, bacterial cells segregate replicated chromosome regions soon after their duplication (with *E. coli* showing some delay), such that sister copies of duplicated chromosome sites are one or more microns apart (43–45). Therefore, during DSB repair occurring away from replication forks, RecA would need to scan thousands if not millions of base pairs within the chromosome(s) for a region that is homologous to that having a break. Interestingly, in *E. coli* and *Caulobacter crescentus* cells, broken sites are moved together, e.g. a cut at one origin regions leads to a transient transversion of the other origin region through the entire cell (28) (in *C. crescentus*, origins are tethered to the cell poles) or in *E. coli*, into the cell centre, where both origins meet (27,29). As opposed to this, in *B. subtilis*, origin sites containing an inducible break site appear to stay within both cell halves during repair via HR (26). A hallmark for RecA activity is the formation of ssDNA-protein filaments *in vitro*, and the arising of filamentous structures *in vivo*, in response to the induction of DNA damage including generation of single DSBs, as well as of replication roadblocks, as has been shown using fluorescence microscopy (26,27,29,30,46). Filamentous forms appear to represent bundles of filaments (27,29) that extend in a time frame of minutes, away from single, induced DSBs, to elongate along the length of rod-shaped cells. While clearly, these structures, termed “threads” in *B. subtilis*, are crucial intermediates during homology search within the chromosome(s), the mode of their dynamic remodelling has been unclear. Using two mutant forms of *B. subtilis* RecA, we show that reduced ATP binding does not strongly alter RecA dynamics *in vivo*, but lack of ATPase activity strongly affects the formation of threads, as well as their dynamic remodelling.

For *E. coli* RecA, it has been proposed that filaments extending along the long axis of the cell search for homology at many sites along their entire length, as has been shown to function *in vitro* (47), to reduce a three-dimensional search for a homologous sequence towards a two-dimensional search. Our finding that ATPase activity is crucial for filament extension and remodelling strongly supports this idea. We show that reduction of ATP binding (and secondary reduction of ATPase activity) has an only mild effect on thread formation by RecA, while loss of ATPase activity is detrimental for this process. Using fast superresolution microscopy analyses, we show that remodelling of RecA filaments occurs within 20 seconds-intervals, and throughout the cell including nucleoids, rather than exclusively along the cell membrane, in agreement with an earlier study (29). At such a speed, RecA filaments could simultaneously test for homology within many chromosome segments; failure to identify sufficient homology would involve ATPase activity to also release RecA-covered ssDNA from duplex DNA, whereby filaments could either shrink when many RecA molecules are unbound and rebound (this can also occur *in vitro*, in the absence of new RecA “loaders”), or simply diffuse vertically along the short axis of the cell, such that eventually, the nucleoid is tested along its length as well as its depth. Three-dimensional diffusion is possible also for long polymers such as dsDNA itself, which shows considerable displacement within a time frame of seconds simply based on Brownian motion (48,49).

We further extended our analyses using single molecule tracking of RecA. We found that trajectories collected from many cells could be well explained assuming three distinct (but interchangeable) populations: we observed a) rapid diffusion throughout the entire cell (Fig. 6), likely representing freely diffusing RecA dimers (purified RecA forms predominantly dimers under physiological salt conditions, Fig. 2A), b) medium-high diffusion, likely due to hopping between DNA strands based on non-specific dsDNA binding by RecA, and c) low-mobility, visible as confined motion on the nucleoids (Fig. 7). Low-mobility is caused by RecA being bound within filamentous structures (Fig. 7), because the induction of a single DSB greatly increased this fraction, at the expense of medium – and high mobile fractions. Of note, most DNA-binding proteins are largely present in constrained motion on the nucleoids, due to non-specific DNA interactions (50), and indeed, a GFP-RecA fusion largely localizes to the nucleoids using epifluorescence microscopy (26). Therefore, the 41% of RecA molecules that we determined may be an overestimate of free RecA diffusion, because the diffusion constant of 0.59 μm^2^ s^-1^ is quite low for a freely diffusing 128 kDa dimeric protein (RecA plus mVenus), suggesting that the high-mobility fraction also contains some (fast diffusing) molecules from the non-specifically DNA-bound fraction. In any event, low mobility of 0.02 μm^2^ s^-1^ obtained in our analyses can only be explained by RecA molecules being part of a very large structure showing extremely low subcellular motion, such as filaments/threads observed in SIM microscopy (Fig. 4). Our data revealing a strong reduction in medium – and fast-moving molecules to be recruited into RecA filaments is in agreement with earlier experiments showing that a redistribution of existing RecA molecules is sufficient for an efficient repair via RecA filaments (51). RecA having reduced ATP-binding ability showed changes in single molecule dynamics similar to those of wild type RecA, in those 30 minutes after induction of a DSB, 40% of RecA molecules were in a low-mobility/filament bound form, from 11% during exponential growth. Because RecA is a highly abundant protein (35), these data suggest that a massive number of RecA molecules is bound to ssDNA even if only a single DNA cut occurs within the genome. Contrarily, ATPase-mutant RecA displayed completely aberrant behaviour: during exponential growth, RecA_K70R_ showed a large, low-mobility population, and less freely diffusive molecules, while induction of a DSBs freed molecules from the low-mobility state. We interpret these findings to suggest that in the absence of ATPase activity, RecA cannot escape from non-productive recombination events occurring during exponential growth. In other word, loss of ATPase activity appears to block disintegration of initial recombination events, such that RecA_K70R_ is erroneously often stuck in low mobility.

**Figure 7.**
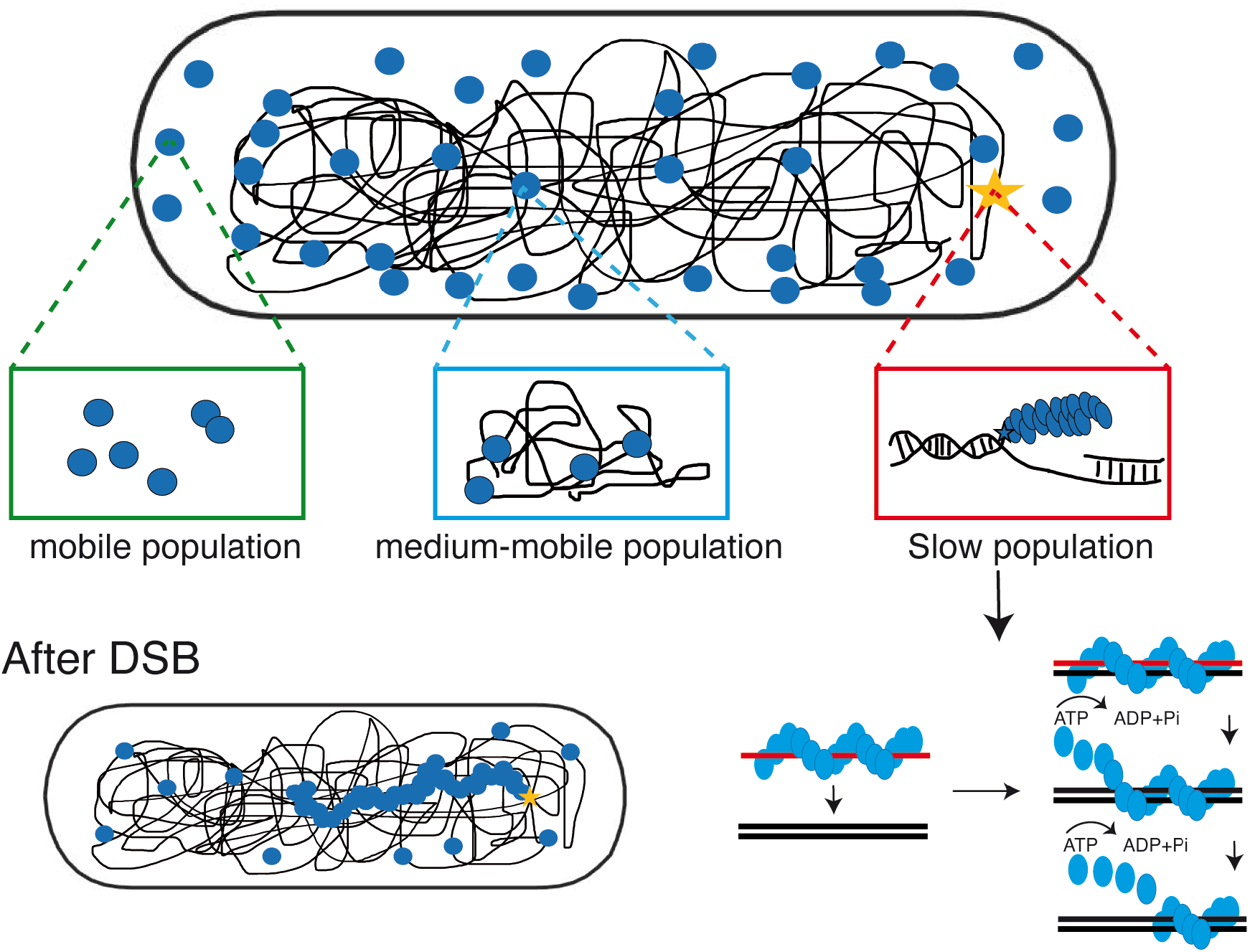
Model for the *in vivo* function of the ATPase-driven formation of dynamic RecA threads. RecA can bind either ssDNA or dsDNA, if it is recruited to ssDNA, e.g., at a site of a DNA break, it forms a helical filament. The helical ssDNA/RecA filament searches for a homologous region of dsDNA. Strand exchange then occurs, and the strand exchange process is driven by product stability, and factors such as RecG and RecU/RuvAB. RecA binding to undamaged dsDNA segments of the genome leads to constrained motion through the nucleoids, and filament formation is achieved by binding to ssDNA from freely mobile and from dsDNA-bound molecules.

In *E. coli*, the intrinsic ATPase activity converts the high affinity binding form, RecA-ATP, to the low affinity binding form, RecA-ADP, driving dissociation from either the hybrid dsDNA products of strand exchange or from undamaged dsDNA. In eukaryotic cells, dissociation of Rad51 or Dmc1 from dsDNA requires the action of a RAD54 family dsDNA-specific translocase which uses the energy of ATP hydrolysis to dissociate the protein from strand exchange products or regions of undamaged dsDNA (17).

Surprisingly, purified ATPase-mutant RecA was defective in nucleating nucleofilaments *in vitro*, in contrast to *E. coli* RecA (52). Because we did observe some filamentous RecA-mVenus structures *in vivo* following DSB induction, we argue that its defect appears to be partially overcome *in vivo*, likely because of the action of RecA loading or accessory factors such as RecO and RarA (37,53). In any event, RecA from *B. subtilis* seems to differ in its molecular properties related to ssDNA binding compared with *E. coli*, so our study also strengthens the important idea that action of RecA at a molecular level, despite being generally conserved between organisms (including Rad51 from eukaryotes), must be viewed considering differing biochemical properties. This agrees with, e.g., RecA from the highly radiation resistant bacterium *Deinococcus radiodurans* operating in a different way during repair via HR than RecA in *E. coli* cells (54). Nevertheless, observations from this study clearly support the perception that the RecA nucleofilaments are tightly regulated by ATPase activity, including both ATP binding and ATP hydrolysis, to remove excessively assembled RecA on ssDNA. This tight regulation is critical to the success of HR *in vivo*.

In conclusion, our work reveals that ATPase mutant RecA retains only little, if any, activity of DNA repair via HR, and that dynamics of filaments/threads in cells is highly perturbed, suggesting that homology search along filaments is a major driver for directional search of sister regions for setting up Holiday junctions. Out data also agree with the idea of ATPase waves within RecA filaments generating a motor-like function that could lead to spatial reorganization of filaments (55). Fig. 7 shows a model in which continued ATPase activity with RecA filaments (only one is shown) leads to homology search along the long axis, to filament extension and diffusion towards the other end of the cell, made possible by the release of non-productive strand exchange events.

While *E. coli* and *C. crescentus* cells set up HJs by joining the DSB site and the sister site, *B. subtilis* cells appear to do this across several micrometres, because break sites do not appear to move, in contrast to directional RecA threads. It will be intriguing to investigate how *B. subtilis* or possibly Gram-positive bacteria in general organize HJ formation in time and space.

## MATERIALS AND METHODS

### Bacterial Strains and Growth Conditions

The bacterial strains and plasmids used in this study are listed in Table S1, and the nucleotides are listed in Table S2. *Escherichia coli* strain XL1-Blue (Stratagene) was used for the construction and propagation of plasmids and *E. coli* strain BL21 Star DE3 (Invitrogen) for the heterologous overexpression of proteins. All *Bacillus subtilis* strains were derived from the wild-type strain BG214. Cells were grown in Luria-Bertani (LB) rich medium at 37°C or 30°C. When needed, antibiotics were added at the following concentrations (in μg/ml): ampicillin, 100; chloramphenicol, 5; spectinomycin, 100; kanamycin, 30. When required, media containing 0.01–0.5% xylose were prepared by adding appropriate volumes of a filter-sterilized solution.

### Construction of strains

RecA, RecA_K70A_ and RecA_K70R_ were visualized as a RecA-mV, RecA_K70A_-mV, RecA_K70R_-mV (for use in Single Molecule Microscopy) or RecA-sfGFP, RecA_K70A_ - sfGFP, RecA_K70R_ -sfGFP (with the goal to use in Structured Illumination Microscopy), fusion proteins expressed at the original locus. The entire *recA* variant ORF’s were integrated into vector pSG1164-mVenus or pSG1164-sfGFP, using *Apa*I and *EcoR*I restriction sites, and BG214 cells were transformed with the resulting constructs, selecting for cm resistance (leading to strains Table S1). This strategy resulted in merodiploid cells: one version was transcribed from the original *recA* promoter, the second was placed under control of the xylose promoter and was not induced. We verified by sequencing that the point mutations were present in the expressed allele. For DSB studies, HO endonuclease system was integrated at *amyE* locus using the plasmid pSG1192, and expression was controlled by xylose addition (0.5% final concentration) (30,56). Transformation of BG214 using was achieved by growing overnight cultures at 30°C and 250 rpm in liquid LB media (10 grams sodium chloride per litter). The next day, we used a 200 ml shaking flask to inoculate 10 ml of freshly prepared liquid modified competence media (MCM) with our overnight culture to yield an optical density of 0.08-0.1 measured at 600 nm (OD_600_). MCM was prepared according to published procedures (57). The prepared culture was grown in MCM at 37°C under constant shaking (200 rpm) to ensure proper aeration till it reached stationary growth phase indicated by an OD_600_ of 1.4–1.6. For subsequent transformation of plasmid or chromosomal DNA, we used an aliquot of 1 ml from that culture and added a total of 1 μg of the respective DNA to it (either plasmid or chromosomal DNA); as a control, we used 1 ml of the same culture without any addition of DNA. Each culture was further incubated at 37 °C in tubes with constant shaking for two more hours, followed by streaking out different amounts of culture aliquots onto fleshly prepared, solid LB-agar plates containing the appropriate antibiotics to maintain selective pressure for the respective strain.

For expression of soluble 6xHis-RecA, 6xHis-RecA_K70A_ and 6xHis-RecA_K70R_ the coding sequence lacking the first 10 codons was amplified by PCR using chromosomal DNA from *B. subtilis* wild type strain BG214. The fragment was further integrated in the expression vector pET28a (Novagen) by *Eco*RI and *Xho*I restriction ligation and brought into the expression host *E. coli* BL21 (DE3) giving rise to the strains pET28a::*recA*HisTag, pET28a::*recA*_K70A_HisTag and pET28a::*recA*_K70R_HisTag.

### Expression and purification of RecA variants

Protein purification was performed in two consecutive steps. The purification of (His)6-RecA, RecA_K70A_ and RecA_K70R_ initially began with affinity chromatography using an ÄKTA Prime apparatus (GE Healthcare) and Nickel-Sepharose columns (HisTrap HP 1 ml, GE Healthcare) and was continued by size-exclusion chromatography using an ÄKTA FPLC apparatus (GE Healthcare) and a gel filtration column (Superdex 75 16/60 GL, GE Healthcare). Prior to purification, the respective proteins were overexpressed in BL21 DE3 cells carrying a pET28a vector (Novagen) with an (indirectly) IPTG-inducible T7 promoter, six encoded histidine and the full gene sequence of the *B. subtilis recA* gene and *recA* variants. Transformants were grown under vigorous shaking in LB-medium at 37°C to exponential phase (OD_600_ 0.6) and induced for 60 minutes with 1 mM IPTG. Subsequently, the cells were centrifuged (20 minutes, 4°C, 5000 rpm) and the pellet was resuspended in HEPES A (50 mM HEPES, 300 mM NaCl, pH 7.5). To prevent protein degradation a protease inhibitor was added (Complete, Roche). Afterwards, the cells were French pressed (AMINCO French Press, Laurier Research Instrumentation) in two consecutive cycles at approximately 20000 psi and the lysate was centrifuged (30 minutes, 4°C, 16000 rpm). The clear supernatant was passed through a filter (pore-size 0.45 μm, Filtropur S, Sarstedt) before injection into the loop of the ÄKTA Prime apparatus (preequilibrated with HEPES A and HEPES B [50 mM HEPES, 300 mM NaCl, 500 mM imidazole, pH 7.5]). The proteins were loaded onto the Nickel-Sepharose column, the column was washed with 20% HEPES B and the protein eluted with 100% HEPES B in fractions of 1 ml and checked by SDS-PAGE. Fractions containing significant amounts of the desired protein were assembled and loaded onto size exclusion chromatography columns (preequilibrated with HEPES A). The peak fractions were analysed by SDS-PAGE and only pure protein fractions were assembled and stored at −80 °C.

### Gel filtration

GF of HisTag-RecA_WT_ and mutants after Nickel-Sepharose columns affinity chromatography continued by size-exclusion chromatography using an ÄKTA FPLC apparatus (GE Healthcare) and a gel filtration column (Superdex 75 16/60 GL, GE Healthcare) standard size is shown with triangles in the upper part of the chromatogram. Fractions in panels showing SDS-PAGE correspond to elution fractions in GF fractions

### ATPase assays

RecA ATPase activity was measured using coupled spectrophotometric enzyme assay. The reaction was performed in 100 μl of assay buffer (6 mmol/l MgCl_2_, 20 mmol/l KCl and 100 mmol/l Tris–HCl, pH 7.4) 1 mmol/l ATP and different concentrations of RecA, RecA_K70A_ and RecA_K70R_ or vehicle (DMSO) and incubated at 37 °C for 3 h. At the end of the incubation, the ATPase activity of RecA, RecA_K70A_ and RecA_K70R_ were assessed by malachite green reagent (Sigma-Aldrich) (0.0812% *w/v* malachite green, 2.32% *w/v* polyvinyl alcohol and 5.72% *w/v* ammonium molybdate in 6 mol/L HCl, and argon water mixed in a ratio of 2:1:1:2, *v/v/v/v*). Reactions were analysed in triplicate at an absorbance of 620 nm. The kinetic analysis of the RecA, RecA_K70A_ and RecA_K70R_ ATPase activity was carried out using a nonlinear regression fit of the experimental points to the Michaelis–Menten equation. Commercially available Salmon Sperm ssDNA was obtained from Sigma-Aldrich.

### Western blotting

*B*. *subtilis* o*r E. coli* cultures (1 ml) were harvested by centrifugation. The pellet was resuspended in lysis buffer (20 mM Tris-HCl [pH 7.0], 10 mM EDTA, 1 mg ml^-1^ lysozyme, 10 g ml^-1^ DNase I, 100 g ml^-1^ RNase I, 1 tablet of Mini EDTA-free, EASY pack (Roche, protease inhibitor cocktail), and incubated for 30 min at 37°C. Proteins were separated by running 12% sodium dodecyl sulphate-polyacrylamide gel electrophoresis (SDS-PAGE) and were transferred onto nitrocellulose membrane followed by blocking with 5% milk in PBST (80 mM Na_2_HPO_4_, 20 mM NaH_2_PO_4_, 100 mM NaCl, 0.2% (v/v) Tween-20). Proteins were probed using a 1:500 dilution (rabbit-α-GFP), 1:1000 dilution (rabbit-α-*recA*) or 1:1500 dilution (mouse-α-His) and secondary antibody was added (goat-α-rabbit-antibody in 1:10000 dilution or goat-α-mouse-antibody 1:10000) after a series of washing steps with PBST. Solution A (100 mM Tris pH 8.5, 2.5 mM Luminol, and 0.4 mM Coumaric acid) and Solution B (100 mM Tris pH 8.5, 0.02% (v/v) H_2_O_2_) were prepared and mixed followed by incubation for 2 min for chemiluminescence detection with ChemiDocTM MP System (BIO-RAD).

### Mass photometry

Oligomeric states of proteins were determined on a One^MP^ mass photometer (Refeyn Ltd., Oxford, UK). Microscope coverslips (1.5 H, 24 x 60 mm, Carl Roth) and CultureWell™ Reusable Gaskets (CW-50R-1.0, 3 x 1 mm, Grace Biolabs) were cleaned with three alternating rinsing steps of ddH_2_O and 100% Isopropanol and dried under a stream of compressed air. Coverslips were coated with Poly-L-lysine by pipetting 7 μl of solution (0.01%, Sigma-Aldrich) between two coverslips, incubation for 30 sec, dipping in and rinsing the coverslips with ddH_2_O after separation, and drying in an air stream. Silicone Gaskets with four cavities were adhered on coverslips. Prior to each measurement, 18 μl of PBS (pH 7.4, RT) solution was pipetted into one cavity and the instrument was focused. 2 μl of protein sample were added, mixed, and measured for 60 s at 100 frames per second using AcquireMP (Refeyn Ltd., v1.2.1). As a mass calibration, NativeMark™ Unstained Protein Standard (Thermo Fisher Scientific) was measured, and data was fit to a linear regression. Proteins were measured either alone or mixed with ssDNA oligonucleotides and 1 mmol/l ATP after incubation for 10 min at 37 °C and concentrations between 50 and 100 nM of RecA_WT_, RecA_K70A_ and RecA_K70R_. All data was analysed using DiscoverMP (Refeyn Ltd., v.1.2.3) (58,59).

### Electron microscopy

RecA filament formation was assayed by incubating 4 μM RecA, RecA_K70A_ and RecA_K70R_ in a buffer containing 25 mM Tris–HCl (pH 7.5), 10 mM MgCl_2_, ATP regeneration system (8 mM phosphocreatine, 10 U/ml creatine phosphokinase), 1 mM DTT, 3 mM ATP, 3% (v/v) glycerol, 7.5 mM NaCl, and 60 nM no fold 90°C ssDNA (generated using primers in Table S2) for 10 min at 37°C. Filaments were stabilized by addition of 3 mM ATPγS and 3 min incubation at 37°C. The samples were spotted onto carbon coated grids (400 mesh) were hydrophilized by glow discharging (PELCO easiGlow, Ted Pella, USA). 5 μl of sample (protein, ATPγS, MgCl_2_, ssDNA) and 1:5 dilutions of the assay were applied onto the hydrophilized grids, respectively, and negatively stained with 2% uranyl acetate after a short washing step with H_2_O_bidest_. Samples were analysed with a JEOL JEM-2100 transmission electron microscope using an acceleration voltage of 120 kV. Images were acquired with a F214 FastScan CCD camera (TVIPS, Gauting).

### Single molecule microscopy and tracking

Cells were spotted on coverslips (25 mm, Menzel) and covered using 1% agarose pads prepared before with fresh S7_50_ minimal medium by sandwiching the agarose between two smaller coverslips (12 mm Marienfeld). All coverslips were cleaned before use by sonication in Hellmanex II solution (1% v/v) for 15 min followed by rinsing in distilled water and a second round of sonication in double distilled water. In contrast to the wide-field illumination used in conventional epifluorescence microscopy, the excitation laser beam used in our setup is directed to underfill the back aperture of the objective lens, generating a concentrated parallel illumination profile at the level of the sample, leading to a strong excitation followed by rapid bleaching of the fluorophores. When only a few unbleached molecules are present, their movement can be tracked. In addition, freshly synthesized and folded fluorophores become visible when the sample is excited again. When an observed molecule is bleached in a single step during the imaging, it is assumed to be a single molecule. Image acquisition was done continuously during laser excitation with the electron-multiplying CCD (EMCCD) camera iXon Ultra (Andor Technology, Belfast, UK). A total of 2,500 frames were taken per movie, with an exposure time of 20 ms (23 frames per second [fps]). The microscope used in the process was an Olympus IX71, with a ×100 objective (UAPON 100×OTIRF; numerical aperture [NA], 1.49; oil immersion). A 514-nm laser diode was used as excitation source, and the band corresponding to the fluorophore was filtered out. Of note, cells continued to grow after imaging, showing that there is little to no photodamage during imaging, while cells stop growing when exposed to blue light (below 480 nm). Acquired streams were loaded into Fiji ImageJ (60). Automated tracking of single molecules was done using the ImageJ plugin MtrackJ (61), or u-track 2.2.0 (62).

### Diffusion analysis of single-molecule tracks

Tracking analysis was done with u-track-2.2.0 (62), which was specifically written for Matlab (MathWorks, Natick, MA, USA). Only trajectories consisting of a minimum of 5 frames were considered tracks and included for further analysis. A widely accepted method to analyse the diffusive behaviour of molecules is by using the mean squared displacement (MSD)-versus-time-lag curve. This provides an estimate of the diffusion coefficient as well as of the kind of motion, e.g., diffusive, sub-diffusive or directed. However, the method requires that within a complete trajectory there be only one type of homogeneous motion and that the trajectory is preferably of infinite length. To distinguish immobile and mobile molecules from each other, we compare the frame-to-frame displacement of all molecules in x and the y directions. Using a Gaussian mixture model (GMM) to fit the probability density distribution function of all frame-to-frame displacements, determine the standard deviations σ_1_ and σ_2_, as well as the percentages F_1_ and F_2_ of the slow and the fast subfractions of molecules, respectively. Finally, the diffusion constants were calculated according 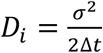, (*i* = 1,2) where *Δt* is the time interval between subsequent imaging frames. Generation of heat maps, analyses of molecule dwell times, and visualization of slow and fast tracks in a standardized cell are based on a custom written Matlab script (SMTracker 2.0) that is available on request (63).

### Structured Illumination Microscopy

Samples at mid-exponential phase were mounted on ultrapure-agarose slides dissolved in LB (1%) for immobilization of cells prior to image acquisition. For localization experiments, image Z-stacks (~100 nm steps) were acquired using brightfield (BF) image acquisition (transmitted light) or structured illumination microscopy (SIM) with a ZEISS ELYRA PS.1 setup (Andor EMCCD camera, 160 nm pixel size; 3× rotations and 5× phases per z-slice; grating period: 42 μm; 100 mW laser line (between 80 and 200 W/cm^2^) at excitation laser wavelength 488 nm; ZEISS alpha Plan-Apochromat 100x/NA 1.46 Oil DIC M27 objective). SIM reconstructions were processed using ZEN-Black software by ZEISS. ImageJ2/Fiji version 1.52p was used for visualization and image processing (60,64–66). Region(s) of interest (ROI) were defined by cell borders using the brush-selection tool to maintain good contrast levels of cellular areas. SIM reconstructions were manually cropped in axial and lateral dimensions, depending on plausibility of cellular positions, using “Duplicate”-function. Signal located outside cell borders was background and was therefore eliminated. Resulting image z-stacks were projected using Fiji implemented “Z-project”-function (e.g., “Average Intensity”), false-coloured and colour-balance adjusted to generate tomographic representations. 3D SIM image z-stacks movies were visualized using Fiji implemented 3D-Project function (with interpolation) for 360° visualization and z-stacks for a tomographic walk-through. Resulting 3D-visualizations were generated with merged channels, processed, and transformed as.avi movies, and finally combined in a sequential manner using Fiji.

## Supporting information

SuppMaterial

## Acknowledgments

This work was supported by the Center for Synthetic Microbiology at the Philipps-Universität Marburg, funded by the LOEWE Program of the state of Hessen, and by DFG-funded research consortium TRR 174.

## Author contributions

R. H.-T. performed all experiments, evaluated data, and co-wrote the manuscript, N.S. performed Mass Photometry assays, T.H. performed Electron Microscopy, G.H. helped with the analysis of Mass Photometry assays and P.L.G. conceived of the study, evaluated data, and co-wrote the manuscript.

## Competing interests

The authors declare no financial of scientific competing interests.

---

**TABLE 1.** Diffusion constants of static, medium-mobile and mobile molecule fractions.

| Strain | # cells | # tracks | D <sup>a</sup> | D<sub>1</sub> <sup>b</sup> | D<sub>2</sub> <sup>c</sup> | D<sub>3</sub> <sup>d</sup> |
| --- | --- | --- | --- | --- | --- | --- |
| **No Damage** |  |  |  |  |  |  |
| RecA<sub>WT</sub>-mV | 138 | 4765 | 0.227 ± 0.018 | 0.036 ± 0.002 | 0.207 ± 0.001 | 0.840 ± 0.002 |
| RecA<sub>K70A</sub>-mV | 126 | 4623 | 0.221 ± 0.015 | 0.036 ± 0.001 | 0.207 ± 0.002 | 0.840 ± 0.001 |
| RecA<sub>K70R</sub>-mV | 132 | 4567 | 0.108 ± 0.017 | 0.036 ± 0.001 | 0.207 ± 0.001 | 0.840 ± 0.002 |
| **After DSB** |  |  |  |  |  |  |
| RecA<sub>WT</sub>-mV | 131 | 4398 | 0.106 ± 0.011 | 0.036 ± 0.001 | 0.207 ± 0.002 | 0.840 ± 0.002 |
| RecA<sub>K70A</sub>-mV | 127 | 4800 | 0.132 ± 0.018 | 0.036 ± 0.002 | 0.207 ± 0.002 | 0.840 ± 0.001 |
| RecA<sub>K70R</sub>-mV | 133 | 5265 | 0.260 ± 0.027 | 0.036 ± 0.001 | 0.207 ± 0.001 | 0.840 ± 0.001 |

995 *aD*, MSD, average diffusion constant of all molecules ( $\mu\text{m}^2\cdot\text{s}^{-1}$ ).996 *bD*<sub>1</sub>, diffusion constant of static fraction ( $\mu\text{m}^2\cdot\text{s}^{-1}$ ).997 *cD*<sub>2</sub>, diffusion constant of medium-mobile fraction ( $\mu\text{m}^2\cdot\text{s}^{-1}$ ).998 *dD*<sub>3</sub>, diffusion constant of mobile fraction ( $\mu\text{m}^2\cdot\text{s}^{-1}$ ).

## Notes

### Competing Interest Statement

The authors have declared no competing interest.

