## Supplementary material for "ATPase activity of *B. subtilis* RecA affects the dynamic formation of RecA filaments at DNA double strand breaks": SuppMaterial

### Supplementary movies

**Movie S1.** SIM reconstruction for $RecA_{WT}$-sfGFP in *B. subtilis*. Cells were grown in the presence of xylose, such that RecA is expressed at physiological levels. Z-stacks resulting from sfGFP channel are merged and projected into tomographic representations. Fluorescent emissions are false coloured in green. Movie Speed 12 fps. 20ms acquisition time, scale bar 2µm.

**Movie S2.** SIM reconstruction for $RecA_{K70A}$-sfGFP in *B. subtilis*. Cells were grown in the presence of xylose, such that $RecA_{K70A}$ is expressed at physiological levels. Z-stacks resulting from sfGFP channel are merged and projected into tomographic representations. Fluorescent emissions are false coloured in green. Movie Speed 12 fps. 20ms acquisition time, scale bar 2µm.

**Movie S3.** SIM reconstruction for $RecA_{K70R}$-sfGFP in *B. subtilis*. Cells were grown in the presence of xylose, such that $RecA_{K70R}$ is expressed at physiological levels. Z-stacks resulting from sfGFP channel are merged and projected into tomographic representations. Fluorescent emissions are false coloured in green. Movie Speed 12 fps. 20ms acquisition time, scale bar 2µm.

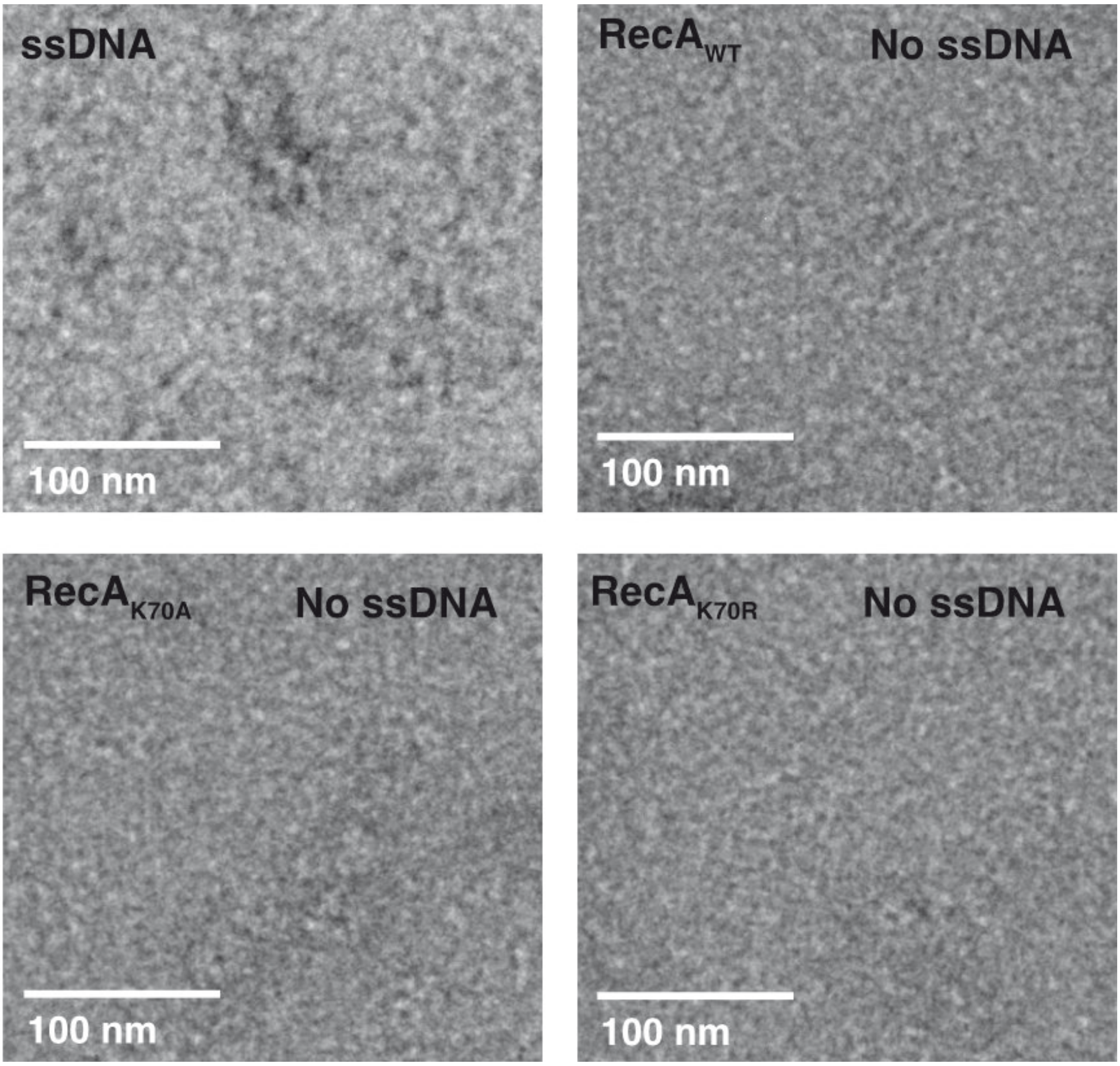


**Fig. S1 Electron microscopy of RecA variant protein with and without ssDNA.** Electron micrographs show filament formation of wild type RecA and no filament formation in RecA mutants without ssDNA.

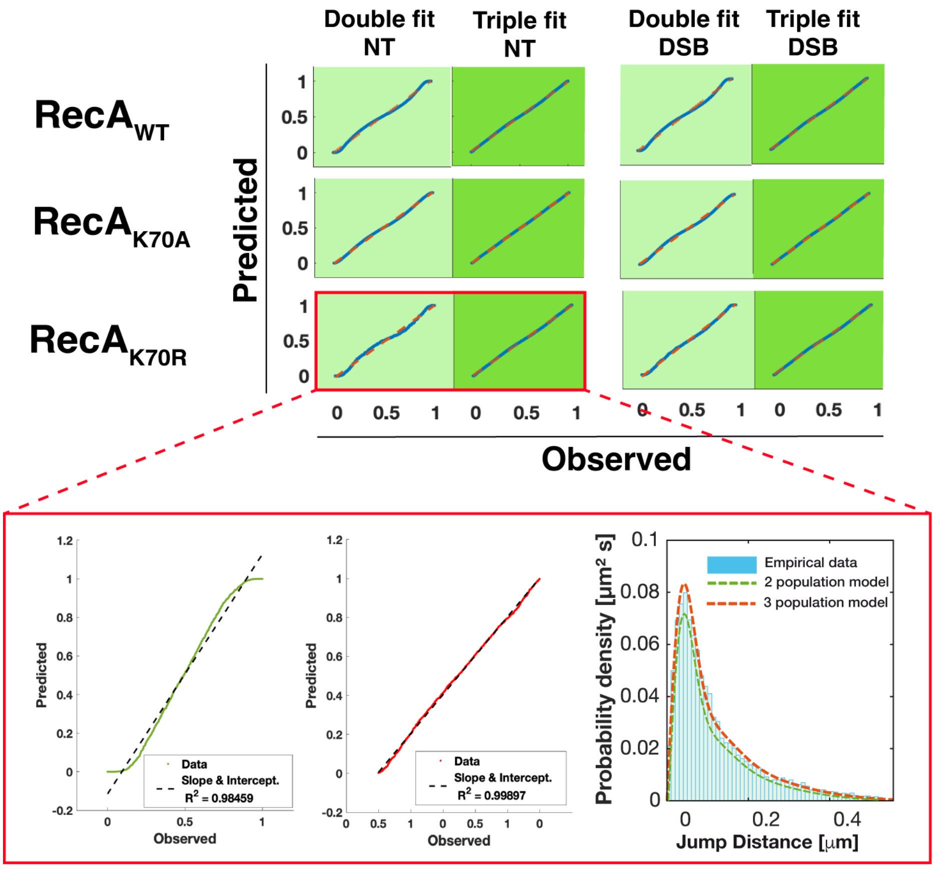


**Fig. S2 Goodness of SQD fit and best model selection.** Probability-probability plots for RecA and every mutant are displayed. In dark green, the model that performs better than the other one, -in light green- in terms of Mean Squared Error (MSE) and R-squared, where using a Triple-population fit clearly performs better, specially at the tails. On the highlighted area, details of the correlation and step-size histogram is shown along with the fit for both models. R-squared coefficients are higher than 0.998 in every of Triple-populations model case.

**TABLE S1** Bacterial Strains and plasmids.

| Strain or Plasmid | Relevant features | Reference or source |
|---|---|---|
| *B. subtilis* | | |
| BG214 | Wild type | |
| BG190 | Δ*recA* *recA::cat* | Alonso et al., 1991 |
| DK52 | *pxyl–HO* endonuclease*::amy -HOcut at spo0J (359°C)* | Kidane et al., 2005 |
| PG7000 | pSG1164::recA$_{WT}$-mVenus$^{cmR}$ / *HO* endo | This study |
| PG7001 | pSG1164::recA$_{K70A}$-mVenus$^{cmR}$/ *HO* endo | This study |
| PG7002 | pSG1164::recA$_{K70R}$-mVenus$^{cmR}$/ *HO* endo | This study |
| PG7003 | pSG1164::recA$_{WT}$-sfGFP$^{cmR}$/ *HO* endo | This study |
| PG7004 | pSG1164::recA$_{K70A}$-sfGFP$^{cmR}$/ *HO* endo | This study |
| PG7005 | pSG1164::recA$_{K70R}$-sfGFP$^{cmR}$/ *HO* endo | This study |
| *E. coli* | | |
| DH5α | *sup*E44 Δ*lac*U169 ϕ80d*lac*ZΔM15 *hsd*R171 *recA*1 *endA*1 *gyrA*96 *thi*-1 *relA1* | New England Biolabs (NEB) |
| BL21 (DE3) | *fhu*A2 [lon] *omp*T gal (λ DE3) [dcm] Δ*hsd*S | New England Biolabs (NEB) |
| pET28a | DH5α pET28a HisTag fusion and expression vector, Kan$^{R}$ | Novagen |
| PG8000 | DH5α pSG1164::recA$_{WT}$-mVenus$^{cmR}$ | This study |
| PG8001 | DH5α pSG1164::recA$_{K70A}$-mVenus$^{cmR}$ | This study |
| PG8002 | DH5α pSG1164::recA$_{K70R}$-mVenus$^{cmR}$ | This study |
| PG8003 | DH5α pSG1164::recA$_{WT}$-sfGFP$^{cmR}$ | This study |
| PG8004 | DH5α pSG1164::recA$_{K70A}$-sfGFP$^{cmR}$ | This study |
| PG8005 | DH5α pSG1164::recA$_{K70R-}$sfGFP$^{cmR}$ | This study |
| PG8006 | BL21 (DE3) pET28a:: recA$_{WT}$-HisTag $^{KanR}$ | This study |
| PG8007 | BL21 (DE3) pET28a:: recA$_{K70A}$-HisTag $^{KanR}$ | This study |
| PG8008 | BL21 (DE3) pET28a:: recA$_{K70R}$-HisTag $^{KanR}$ | This study |

**TABLE S2** Oligonucleotides used in this work.

| Name | Sequence[a,b] |
|---|---|
| RecAamydw310 | *TAT*GAATTCACCCCCTTCTTCAAATTCGAGTTCTTCTTG |
| RecAamyup315 | *GCA*GGGCCCATGAGTGATCGTCAGGCAGC |
| Hoendoup304 | *CA*GGGCCCAGGAGGTTACCGAATGCTTTCTGAAAACACGAC |
| Hoendodw305 | *TAT*GAATTCTTAGCAGATGCGCGCACC |
| mRFPup | *CAG*GGTACCATGGCCTCCTCCGAGGAC |
| mRFPdw | *TAC*GGGCCCCCCCACCGGGCGCCGGTGGAGTG |
| RecAORFdw | *AA*GGGCCCTTCTTCAAATTCGAGTTCTTCTTGTG |
| RecAORFup | *AC*GAATTCATGAGTGATCGTCAGGCAGCCTTAGATAT |

[a] Non-encoded bases introduced as clamps are shown in italics. Restriction sites are underlined.

[b] The location is indicated by the first 5' nucleotide and the replicon where the sequence is located. Accession numbers are *recA* (CP053102.1 (1764677 to 1765723) of *B. subtilis.*
